# Value computations underlying human proposer behaviour in the Ultimatum Game

**DOI:** 10.1101/100313

**Authors:** Erdem Pulcu, Masahiko Haruno

## Abstract

Interacting with others to decide how finite resources should be allocated between parties which may have competing interests is an important part of social life. Considering that not all of our proposals to others are always accepted, the outcomes of such social interactions are, by their nature, probabilistic and risky. Here, we highlight cognitive processes related to value computations in human social interactions, based on mathematical modelling of the proposer behavior in the Ultimatum Game. Our results suggest that the perception of risk is an overarching process across non-social and social decision-making, whereas nonlinear weighting of others’ acceptance probabilities is unique to social interactions in which others’ valuation processes needs to be inferred. Despite the complexity of social decision-making, human participants make near-optimal decisions by dynamically adjusting their decision parameters to the changing social value orientation of their opponents through influence by multidimensional inferences they make about those opponents (e.g. how prosocial they think their opponent is relative to themselves).

## Authors’ Significance Statement

Humans are capable of developing sophisticated strategies for negotiating how finite resources should be distributed between parties with competing interests. This study describes a cognitive model implementing value computations in risky and uncertain situations, where one’s terms may be accepted or rejected depending on how others value them. Surprisingly, despite its everyday and socio-political importance, the evaluation of risk and uncertainty in human social interactions that involve the distribution of monetary resources has not previously been studied using a computational framework. In an ecologically valid experimental design, we provide quantitative evidence to suggest that people make nearly optimal decisions in social interactions, as they would in a non-social value-based decision-making context, and that these decisions are influenced by the human ability to dynamically adjust the decision parameters, particularly those that depend on how the individual represents different dimensions of the opponents’ social value orientation.

## Introduction

Using the optimal decision-making strategy in non-social and social contexts is a key challenge of our everyday life, and from a broader perspective it is closely linked to our survival. Social contexts demanding optimal decision-making also involve negotiating with others over terms which might result in fair or unfair resource distributions (1–4). A common psychological observation would suggest that the optimal strategy in such social interactions (such as the Ultimatum Game(1, 3), or the Prisoner’s Dilemma(5)) should consider how others will perceive the extent of our cooperative/competitive intensions. This is a task that requires simulating others’ valuation processes. However, because our knowledge of other people’s state of mind is limited, we might miscalculate their reactions. As a result, the proposals that we make are not always accepted. In other words, the outcomes of social interactions which involve the distribution of resources between two parties are inherently probabilistic and risky. This implies that there should be a degree of overlap between cognitive models which account for economic decision-making under uncertainty (6) and those which can capture human behaviour in social interactions. However, there is limited work on computational models of value-based decision-making in social contexts, and it is not known whether similar approaches are used across nonsocial and social situations.

Previous theoretical work demonstrated that it is possible to sustain mutual cooperation even when resources are distributed unfairly between two individuals (7). Using the example of a simple social economic game (i.e. Prisoner’s Dilemma), Press and Dyson demonstrated that in order to sustain mutual cooperation under unfair conditions, the player who aims to establish favourable terms for himself still needs to give enough incentive to his opponent. Thus, the player needs to have good understanding of the opponent’s underlying value function to predict at which stage the opponent might change her strategy and stop cooperating. In this context, human social interactions have an intrinsic element of risk and uncertainty; one side setting the terms of interpersonal cooperation should consider the other side’s rejection possibility (i.e. proposer behaviour in the Ultimatum Game). Additionally, with any move that the player makes towards maximising his own payoff by offering conditions that are not in harmony with the valuation of the opponent, the player risks entering a domain where the opponent’s rejection probability increases. These conditions highlight a social interaction scenario in which a player who is interested in maximizing his payoff needs to make value-based decisions while incorporating his opponent’s rejection probability (e.g. the Balloon Analogue Task(8) in value-based domains; or the proposer behaviour in the Ultimatum Game(3) in social decision-making domains). However, value computations in such social interactions have not been studied experimentally or quantitatively.

Here, we designed a behavioural experiment to capture these decision-making processes. We focused on human participants’ Ultimatum giving behaviour and used this common behavioural economic measure as an experimental probe of how humans tackle resource distribution problems. Our primary aim was to construct a formal cognitive model that can account for how others’ valuation processes are integrated to self-decision values during social decision-making. To do so, we tested the degree to which participants utilise computational models that integrate outcome probabilities and reward magnitudes into expected values. In the context of acting as proposers in the Ultimatum Game, expected value computations would require integrating the inferred acceptance probability of one’s opponent with potential self-reward magnitudes (see Materials and Methods for mathematical definitions). Although computational models of decision-making under uncertainty are relatively well established (9), it is not known how well social decision-making models with a comparable structure can account for human behaviour during social interactions.

In order capture the necessary components of a value-based decision-making model experimentally (i.e. outcome probabilities and reward magnitudes), we asked participants to: (i) learn the underlying value functions of two distinct computerised agents with different Social Value Orientations (SVOs^1,2^; one prosocial, the other individualistic; categorically defined with respect to their degree of prosociality) by observing their Ultimatum acceptance preferences (see Fig. 1 and legends for the experimental design); and (ii) transfer this information to make value-based decisions between one of two Ultimatum offers to be given to agents whom they have observed in the learning sessions. Furthermore, in order to test the prediction that value computations may be modulated differently across non-social and social contexts, we used a probabilistic value-based risk decision-making paradigm as a control condition (Fig. 1). Our *a priori* prediction was that, unlike a stable preference usually seen in non-social contexts, people exhibit a dynamic adjustment in social contexts, which allows them to adapt their behavioural strategies to the changing characteristics of their opponents.

**Fig. 1.**
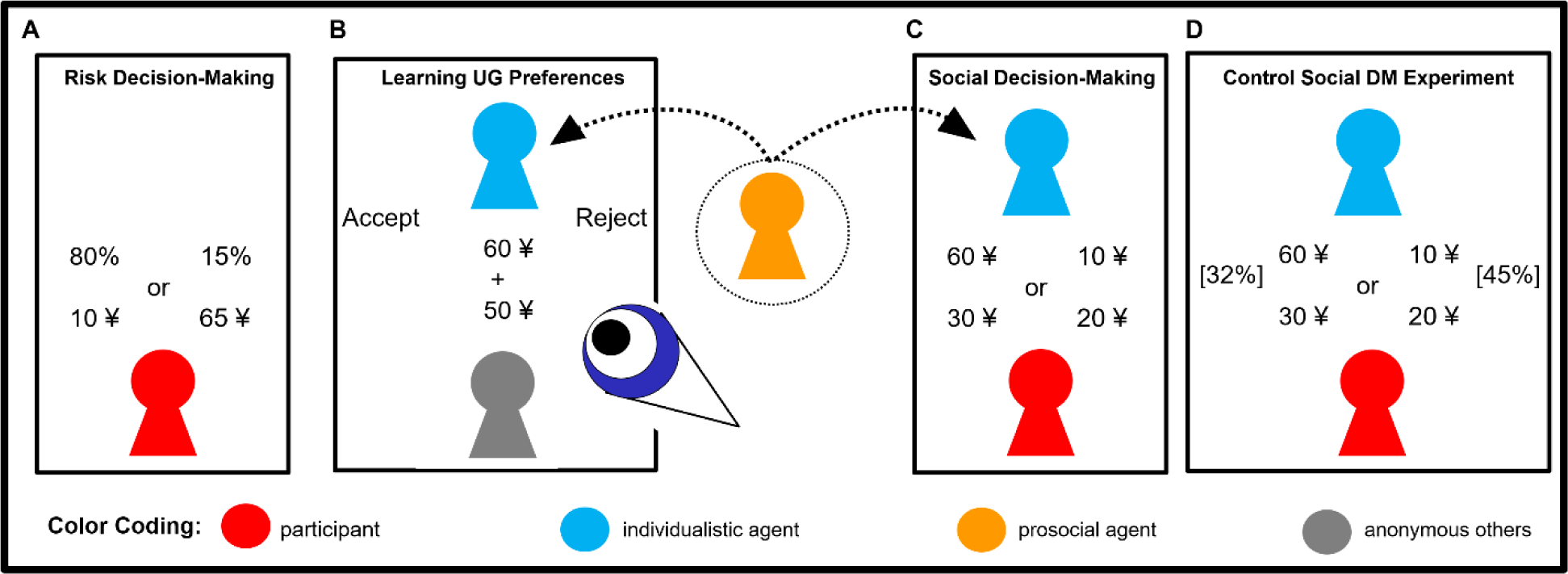
Details of the experimental paradigms. In total participants completed 700 trials: 100 trials in task A, 360 trials in task B and 240 trials in task C. (A) Outline of the baseline risk decision-making task, where participants’ risk perception in a nonsocial context was evaluated. (B) Participants completed two observational social-learning sessions (represented by the schematic blue eye observing the Ultimatum Game interaction), where they were asked to predict the Ultimatum acceptance preferences of two social agents with different Social Value Orientations (SVO) who were responding to offers coming from different anonymous individuals, blue, individualistic; orange, prosocial agent (colour coding is consistent in all subsequent figures). (C) Following the observational social-learning sessions, participants completed a social decision-making experiment in which they were asked to give Ultimatum offers to those social agents from a binary selection. (D) Outline of the control social decision-making task administered to an independent cohort, in which the social agents’ acceptance probabilities (shown inside square brackets) were given explicitly alongside the Ultimatum offers. All tasks were self-paced. UG, Ultimatum Game; DM, decision-making.

## Results

**Social Learning Session.** Participants were able to predict the Ultimatum acceptance preferences of social agents with different SVOs ~ 70% correctly. Prediction accuracy was significantly higher than random guessing for both social agents (Fig. 2A; t-tests from 0.5; all t>22; pcO.OOl, Bonferroni corrected). Participants were able to predict the decisions of the prosocial agent significantly more frequently than the individualistic agent (t=-9.94; p<0.001). Participants’ prediction accuracy closely followed the subjective valuation of the social agents: predictive accuracy was low around the indifference point of the agents’ subjective valuations (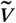 = 0) and increased whenever the subjective valuation of the agents was either very negative or positive (Fig. 2B).

**Fig. 2.**
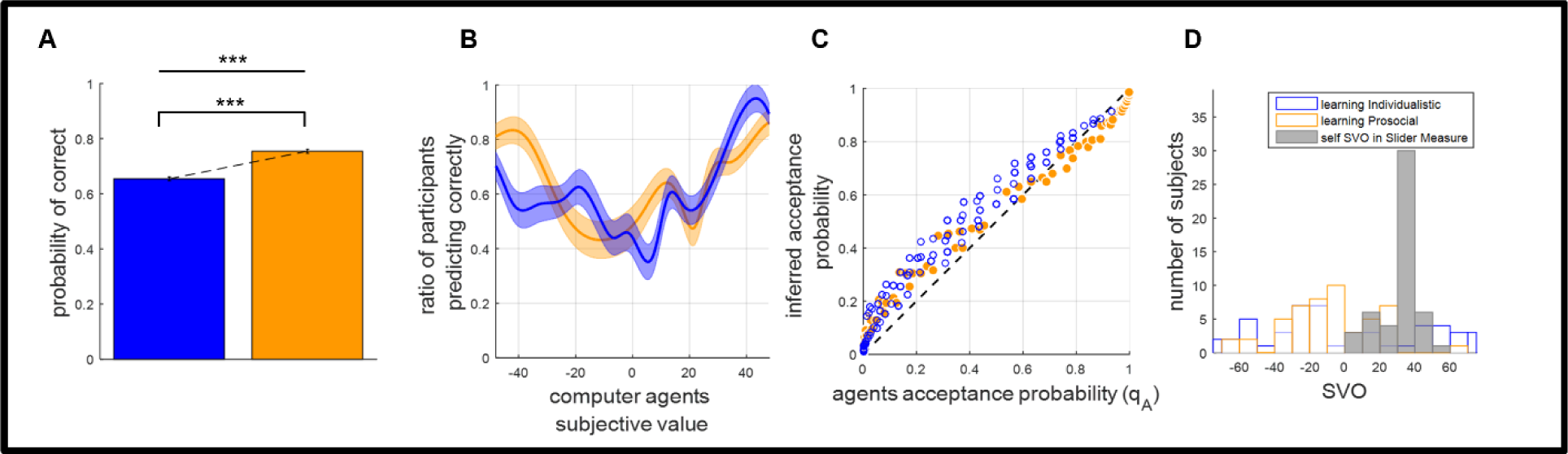
Summary of results from the observational social-learning session. (A) Participants’ prediction accuracy was significantly higher than random guessing and higher for the prosocial agent relative to the individualistic agent (*** P<0.001, error bars show SEM). (B) Prediction accuracy followed the subjective valuation of the social agents, irrespective of their SVOs. (C) Encoded value functions were highly and significantly correlated with the social agents’ actual value functions (r>.97, P<0.001), and had non-linear properties. (D) Participants SVOs calculated on the basis of their decisions in the learning sessions had significantly different distributions than their SVOs measured by the Triple Dominance Measure (*D*>0.60, P<0.001), suggesting participants actively made decisions which are not in accordance with their SVOs to learn about these social agents.

The decisions of the computerised social agents were generated by a model which was derived from the SVO framework ((1); also see Materials and Methods). We fitted the same generative model to the participants’ predictions in the learning session to estimate the inference of their opponents’ value function 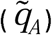. The encoded value functions estimated from the learning session 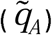 correlated highly and significantly with the agents’ actual value functions (*q*_*A*_; all r(49)>.97, all p<0.001, Bonferroni corrected, see Fig. 2C). This learning model had significantly better fitting relative to a model that has the same number of parameters but makes random predictions in terms of -log likelihood values (t-test from 0.69; all t<-81, all p<0.001, Bonferroni corrected) for both social agents. The pseudo-R^2^ values (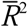; adjusted for the sample size and the number of free parameters(10,11)) of the model were 0.233 and 0.378 for the individualistic and the prosocial agents, respectively. McFadden (1974) suggests that 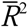 values between 0.20 and 0.40 indicate highly desirable model fitting. Plots of the encoded value functions against the actual value functions revealed non-linear properties (Fig. 2C). The best fitting learning model also had significantly better fitting relative to two alternative reinforcement learning models which were fitted to the participants’ behaviour in the learning sessions (see Materials and Methods, all F_2,147_>168, all p<0.001, Bonferroni corrected). The model also predicted human behaviour significantly better relative to a Bayesian Ideal Observer model that can track social agents’ acceptance probabilities over a numerical grid of self and other reward magnitudes (mean predictive accuracy of the Ideal Observer model across both learning sessions: 78.9% vs 63.3%; main effect of learning model, F(1,98)=283.59, p<.001).

A comparison between participants’ SVO in terms of degrees, as measured by the SVO Slider Measure(2) and based on participants’ choices in the learning sessions, suggested that participants should be making predictions actively to encode the preferences of the computerised agents (i.e. not choosing for oneself; 2-sample Kolmogorov-Smirnov tests, all *D*>0.62; all p<0.001, Bonferroni corrected; see Fig. 2D). Similarly, the distribution of participants’ SVO based on participants’ predictions in the learning sessions was significantly different between the individualistic and prosocial agents (*D*=0.28; p=0.03), suggesting that participants relied on different predictions to learn about their opponents’ underlying value functions.

We were also interested in understanding the participants’ affective reactions to these social agents. In order to address this issue, after each learning session, we asked the participants to rate the imagined personalities of these social agents on a number of different domains. The domains were related to social constructs such as the SVOs of the social agents and how much the participants would like the agent in real life. Responses to these questions (please see Fig. S1A legends) showed that the prosocial agent was rated consistently higher relative to the individualistic agent, which conforms to the general intuition that prosocial individuals would be regarded more positively in real life (2×4 multivariate ANOVA showing main effect of agent F_3,294_ =3.01, p=0.03 and main effect of the interaction term F_3,294_ =10.163, p<0.001; see Fig. S1A). Particularly the participants’ responses to Q3, in which we asked how many people they know in real-life who behave similarly to the computerised agents whose decisions they observed, suggests that our experimental manipulation successfully mimicked interactions with real human opponents (1-sample t-test relative to 0 (i.e. computerised agents’ decisions do not resemble any people the participant knows); all t>13; all p<.001, Bonferroni corrected).

**Value-based Decision-Making.** All participants completed a value-based risk decision-making experiment in which they were asked to choose between 2 probabilistic gambles (see Fig 1A for the task screen). We used this experiment as our control condition to evaluate the degree to which value-based decision-making models account for human behaviour in both non-social and social settings. Model selection based on group-wise sum of BIC (Bayesian Information Criterion) scores suggested that the best fitting model to participant choices was the one with a power utility parameter that modulates the reward magnitudes and integrates the magnitudes with outcome probabilities to compute the expected value difference between available options (see Model 3 in Supplementary Materials and Methods for mathematical descriptions). The best fitting model in the value-based risk decision-making experiment had a -log likelihood value of 0.242/trial and group-wise sum of BIC score of 2699.

We also considered that accumulated winnings over time might have an influence on participants’ choice behaviour. To evaluate this possibility, we included an additional free-parameter to the best fitting model to account for the influence of accumulated wealth down the trials. However, this model did not improve the model fits any further, and the value of the added free-parameter (linearly scaling accumulated winnings) approached zero (mean±SD=1.682×10^−4^ ± 5.056×10^−4^), suggesting accumulated winnings may not have profound influence on participants’ choice behaviour.

**Social Decision-Making.** Upon completion of the learning sessions, our participants progressed with the social decision-making experiment(s), where they interacted with these social agents by making Ultimatum offers for 120 trials against each opponent (see Fig. 1C for task screen).

After each social decision-making block, participants were asked to rate how much weight they put on other’s inferred acceptance probability and/or their self-reward magnitudes while making decisions (on a scale from 0 to 10). Here, a rating of 0 would refer to making decisions only based on inferred acceptance probabilities; a rating of 10 would refer to relying solely on self-reward magnitudes; and a rating of 5 would mean their equally weighted integration. In accordance with our predicted value-based social decision-making model (i.e. making offers based on the expected value difference), our participants reported that they considered both the other’s inferred acceptance probabilities 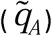 and how much they would win if their offer is accepted (i.e. self-reported integration weights were significantly different than relying on either self-reward magnitudes or encoded acceptance probabilities; all p<0.001, Bonferroni corrected, see Fig. 3A). However, the self-reported integration of these decision variables while making the offers was significantly different than an integration with equal weighting (i.e. participants reported putting more weight on inferred acceptance probability 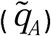, all t>10.46; all p<0.001, Bonferroni corrected). Furthermore, there were no significant differences between the integration weights reported for the prosocial and the individualistic agents, suggesting that our proposed value-based social decision-making model should apply irrespective of the opponents’ SVO (also in Fig. 3A).

**Fig. 3.**
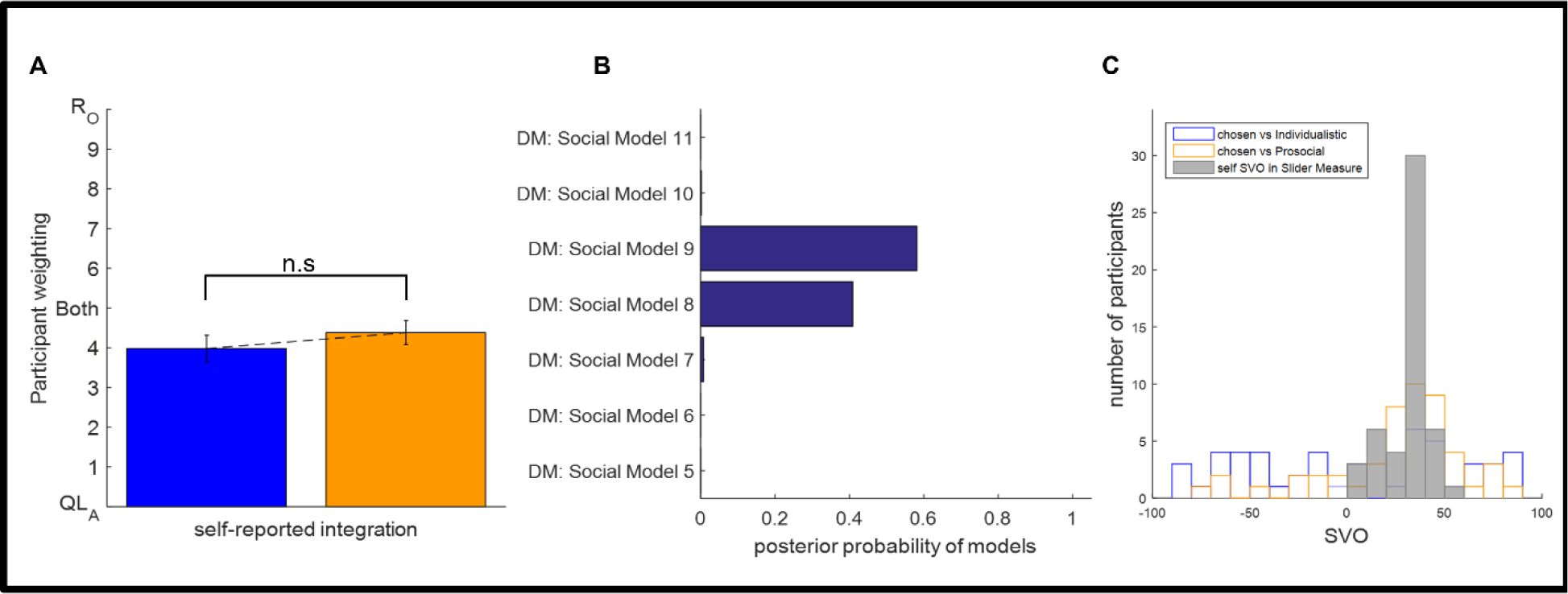
Participants’ self reported value integration and model selection in the social decision-making experiment. (A) In line with our proposed model, participants reported that they considered both their self-reward magnitudes and others’ inferred acceptance probabilities. Mean of bars are highly and significantly different than 0 (only consider inferred acceptance probability, i.e. QL_A_) and 10 (only consider self-reward magnitude, i.e. R_o_; all P<0.001). Self-reported integration weights were comparable against the prosocial and the individualistic agents (n.s: not significant, error bars show ±SEM). (B) Model selection based on Bayesian posterior probability weights recommends Model 9 (longer bars indicate better fitting), which computes the expected value difference between the offers by making use of the power utility and the probability weighting function. (C) Participants’ SVOs calculated on the basis of their decisions in the Ultimatum giving experiments were distributed significantly differently than the distribution of their actual SVOs in real-life (all *D*>0.26; all P≤0.056). The vectors of the SVOs were calculated based on the chosen options and did not correlate with the participants’ actual SVO as assessed by the Slider Measure (all P>0.32).

Model fits (see Supplementary Materials and Methods for mathematical descriptions) revealed that the best fitting social decision-making model was the one which modulated the other’s inferred acceptance probabilities 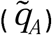 nonlinearly by a probability weighting function and integrated this information with self-reward magnitudes modulated by a power utility parameter that captures participants’ risk attitude (i.e. Model 9; see Fig. 3B). In Social Decision-Making Model 10, we used participants’ self-reported integration weights (as in Fig. 3A) as an additional parameter to allow unequally weighted integration of the perceived probabilities and reward magnitudes. Bayesian posterior probability weights based on the group-wise sum of BIC scores suggested that Model 10, along with others which mainly rely on either reward magnitudes or inferred acceptance probabilities could not account for the participants’ choice behaviour. A complementary Bayesian Model Selection approach also favoured the best fitting model (i.e. Model 9), which had greater exceedance probability than the second-best model (0.73 vs 0.19).

One might argue that our approach to modelling participants’ choices in the social decision-making task neglects any learning which might happen in parallel during this stage. If learning continues to take place during the social decision-making period, the predictive accuracy of our model, which considers the inferred acceptance probability 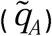 solely based on the learning session, should gradually decay down with increasing trials. In order to investigate this possibility, we segmented the trials in the social decision-making task into three temporal sections: early (1–40); middle (41–80); and late (81–120) trials and compared the predictive accuracy of our model across these sections. This control analysis showed that model predictions were highly stable (all F_2,147_ <0.31; all p > 0.73), suggesting that any additional learning which might take place during the decision-making period would not have a profound effect on the inferred acceptance probabilities participants used to compute the expected value of offers.

Further analysis done by comparing participants’ SVOs calculated on the basis of the offers participants made in the Ultimatum Game versus their SVOs measured by the Slider Measure suggested that the SVOs were distributed significantly differently (Kolmogorov-Smirnov tests; all *D*>0.26, p≤ 0.056, uncorrected; see Fig. 3C). These numerical values (i.e. the SVOs calculated based on Ultimatum offers versus the values from the Slider Measure) were not correlated either (-.141< r (49)<−.035, 0.32< p <0.81). In tandem, these results limit the possibility that, in the social decision-making experiments, participants performed using other cognitive models related to the SVO framework that we did not consider; and lend further support to our prediction that participants used a cognitive model with a structure similar to value-based decision-making under risk and uncertainty.

**Modulation of value-based decision-making in non-social and social settings.** So far, the main difference between non-social and social decision-making experiments is that during social decisionmaking, people are engaged with additional cognitive processes which involve nonlinear probability weighting to compute the expected value difference between the options they face (i.e. the differences between Models 3 (non-social) and 9 (social)).

Although inferred probabilities used in the social decision-making task reflects the true nature of social interactions in everyday life (i.e. one can never know the exact numerical values of the other’s acceptance probabilities), it is important to point out that there were structural differences between the value-based risk decision-making experiment, where the probabilities were given explicitly, and the social decision-making experiment, where the probabilities were experience-based (i.e. inferred). In order to understand the degree to which such differences determined the best fitting model, we conducted an additional control experiment in an independent cohort (n=19; 63.2% males; [mean±SD] age: 21.3±1.9; SVO:27.8±15.1), in which participants completed all value-based decision-making and social-learning tasks as before, but were shown the opponents’ acceptance probabilities explicitly in the social decision-making sessions (Fig 1D). The number of participants were relatively lower in the control experiment. Nevertheless, the effect sizes to detect between-group differences were still high (based on the simple behavioural results reported in Fig 2A; main experiment Cohen’s *d*=1.41; control experiment Cohen’s *d* =1.22).

The explicit presentation of others’ acceptance probabilities led to a number of differences. First of all, participants were able to accumulate higher monetary earnings in this condition (main effect of explicit probabilities (F(1,67)=3.231, p=.077), indicating a marginally more optimal, but not statistically significant, expected value difference computation. As one might expect, the best fitting model was analogous to the one which accounts for the best choice behaviour by the participants in the non-social value-based decision-making experiment (i.e. integrating explicit probabilities with reward magnitudes modulated by a power utility parameter). This means that value computations which involve nonlinear probability weighting are unique to social interactions in which the others’ valuation processes are inferred, whereas power utility modulation of reward magnitudes accounting for people’s risk attitudes is an overarching process across non-social and social contexts.

In the next step we wanted to further decompose the differences between value-based decision-making in non-social and social contexts. To begin with, we conducted a model-free analysis of the participant’s choices by analysing the proportion of risky decisions made in each domain, focusing on the trials where participants were asked to choose between low probability-high magnitude and low magnitude-high probability options. This analysis revealed that the frequency of risky choices, after controlling for the number of trials meeting the criteria described above, was not significantly different across non-social and social domains (Fig. 4A; F_2,147_=3.263, p=0.073, Bayes factor for group differences BF_10_=0.516).

**Fig. 4.**
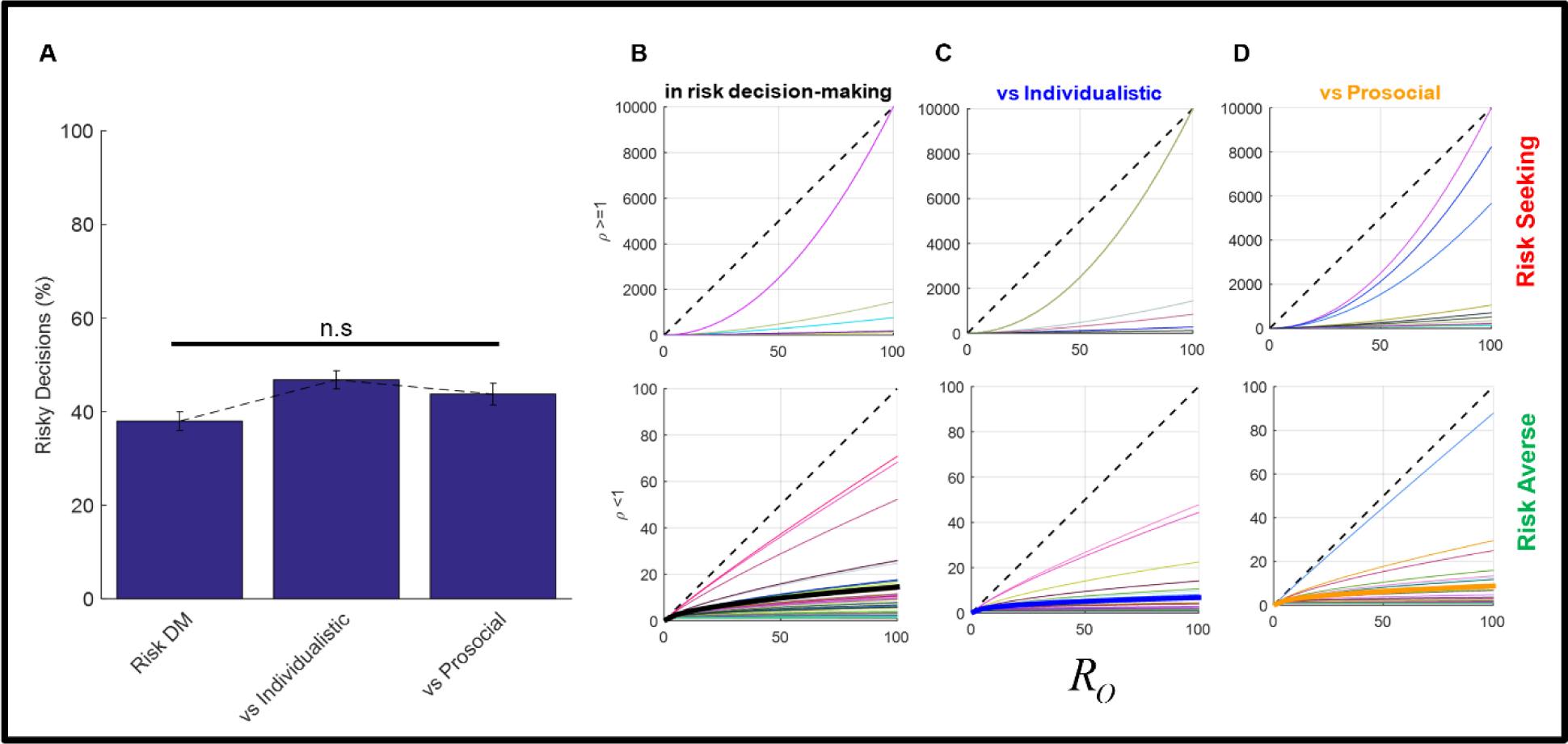
Risk decision-making across non-social and social domains. (A) Proportion of risky decisions made in each domain was not significantly different (P=0.07). Parameter estimates modulating the participants gains (*R_o_*) in each domain: (B) value-based risk decision-making; (C) Ultimatum giving against the Individualistic opponent; (D) Ultimatum giving against the Prosocial opponent. In panels B~D, upper panels show the power utility curves for risk-seeking preferences, whereas the lower panels show risk-averse preferences. Thin lines with the same [R,G,B] colour coding specify each subject’s power utility curve, and thick lines in black, blue and orange colours show the population means for each domain, where we observed a pronounced risk aversion across all domains (P<.001, Bonferroni corrected).

By estimating the power utility parameters separately for the value-based risk decision-making experiment (i.e. the non-social context) and the two Ultimatum giving experiments against different social agents (i.e. social context), we were able to show that on average our participants displayed a pronounced risk aversion across all domains (all p<0.001, Bonferroni corrected relative to risk neutrality *ρ* = 1, see Fig. 4B-D for power utility curves). These parameters were estimated separately in each domain and correlated significantly with the proportion of risky decisions the participants made in those domains (all r(49) >.421, all p<.0024, Bonferroni corrected for pairwise comparisons). Although our participants were slightly more risk-seeking against the individualistic agent, this difference was not significantly higher (p=0.638, see Fig. S2), supporting our *a priori* prediction at the population level that people should not be acting in a consistently risk-seeking manner against one type of social agent.

The power utility parameter estimates from the non-social value-based experiment were not significantly correlated with the estimates from the social decision-making experiments (all r(49)<.27, all p>.06, Bonferroni corrected). Furthermore, power utility parameters were not correlated within the social decision-making domain either (r(49)=.20, p=.173), indicating that people may have an independent risk attitude in social interactions. However, in sharp contrast to these findings, the risk parameters were significantly correlated when the others’ acceptance probabilities were given explicitly (r(18)=.55, p=.015).

A mostly comparable picture emerged for the probability weighting parameters estimated from the Ultimatum giving experiments (Fig 5). Despite individual variability, parameter estimates were not significantly different than ⟀=1 at the population level (i.e. where ⟀=1 describes the diagonal line where the actual and perceived probabilities are equal; all p>.06) and the population mean of these estimates were comparable between individualistic and prosocial agents (p>.91). However, within-subjects correlation for the probability weighting parameters was not significant (r(49)=−.065, p=.65), indicating that probability weighting in social decision-making is not a hardwired trait applicable to different scenarios, but rather adaptive to the changing characteristics of one’s opponents.

**Fig. 5.**
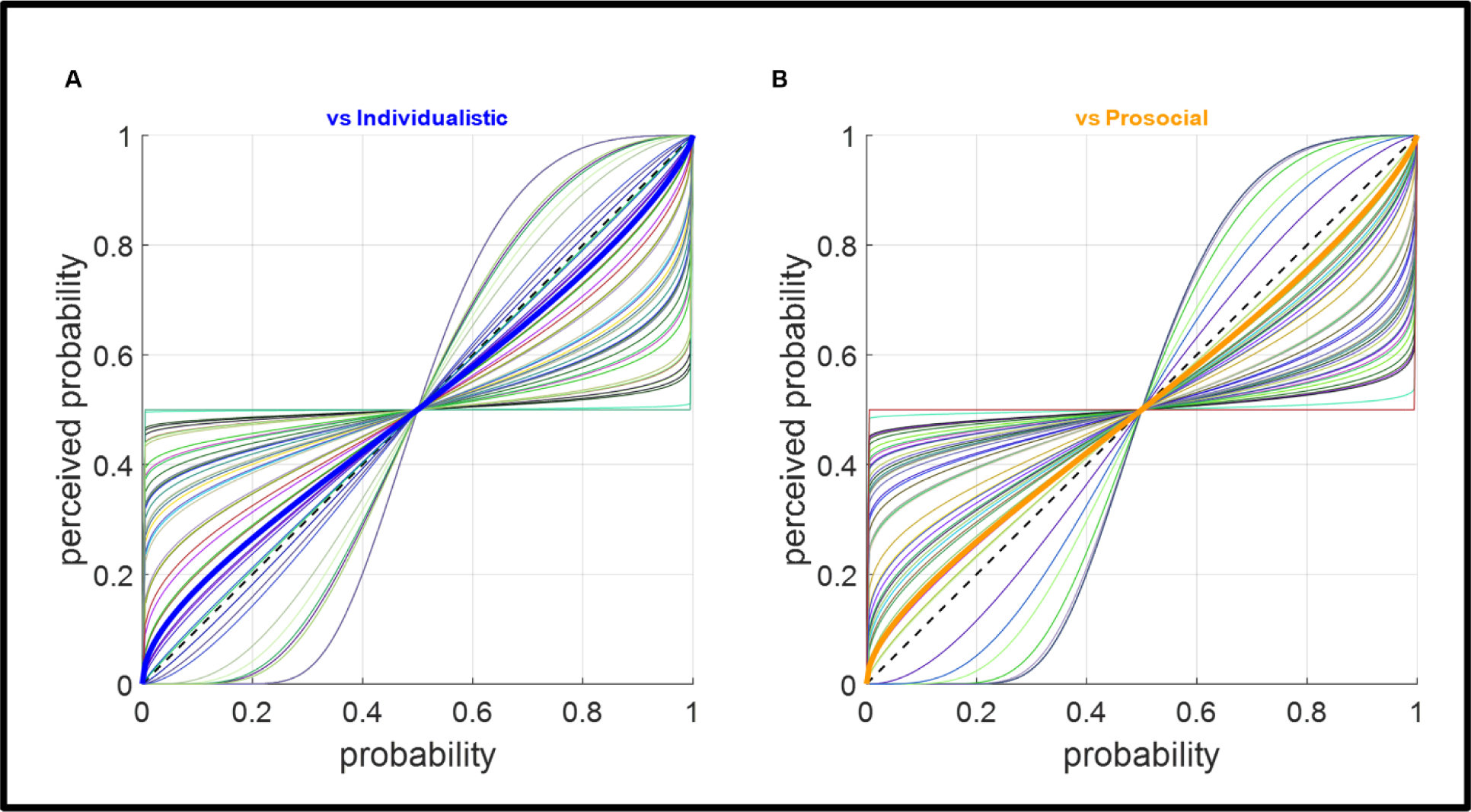
Probability weighting in social decision-making (based on log_2_ functional form). At the population level, parameter estimates (thick blue and orange lines showing the population mean) were not significantly different than 1 (diagonal dashed lines where the actual and perceived probabilities are equal; all p>.06), and these estimates were statistically comparable between individualistic and prosocial agents (p>.91). In both panels, thin lines with the same [R,G,B] colour coding specify each participants’s probability weighting curve.

Intriguingly, the individual variability observed for the power utility and the probability weighting parameters mostly converge at the population level, and choice probability curves suggest that human participants make nearly optimal decisions in giving Ultimatum offers as they make non-social value-based decisions under uncertainty (Fig. 6).

**Fig. 6.**
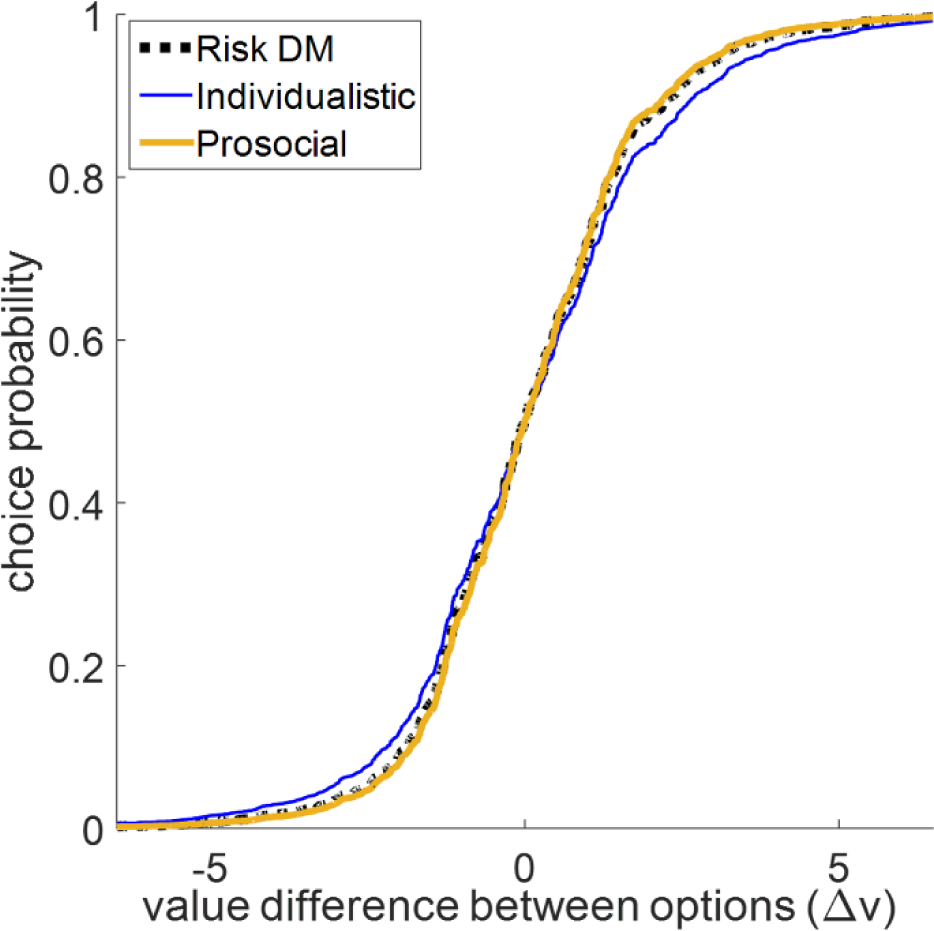
Choice probability curves across non-social and social decision-making shows that human participants can make nearly optimal decisions while giving Ultimatum offers as they would in a non-social context (i.e. value-based risk decision-making. Fig 1A). X-Axis shows the value difference between each option computed according to Model 1 (Eq. 5).

The preceding analyses of the parameter estimates and choice probability curves suggested that human participants show a degree of adaptation during social decision-making which allows them to make nearly optimal decisions as they would in a non-social setting. Next, we wanted to explain the “social” risk and probability weighting parameters estimated from our Ultimatum giving experiments by a number of predictive variables to evaluate the extent to which people’s decision parameters are influenced by social variables describing degrees of prosociality. Our hypothesis was that people’s decision parameters in social interactions should depend on their SVO; their inference about the SVO of their opponents (*SṼO*; including one’s uncertainty about this estimate); and how prosocial they think their opponent is relative to themselves. Here, the relative prosociality term (i.e. difference between self and other) allows us to model the extent of the parameter adjustment in social interactions, particularly in situations where people judge their opponents to be more or less prosocial than themselves. To test this prediction, we constructed two multiple linear regression models with these predictive variables:

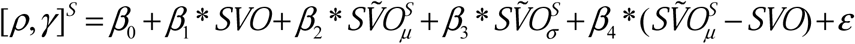

where [*ρ, γ*]^*S*^ are the risk and the probability weighting parameters, respectively, estimated separately from the Ultimatum giving experiments, and *S* ϵ {*i, p*} defines whetherthe opponent is individualistic or prosocial. Here, it is important to point out that participants’ inferences about their opponent’s SVO (*SṼO*^*s*^) depend on their encoded value function 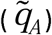 from the learning sessions, which has a stochastic nature (i.e. a sigmoid function). In order to get the best estimate of the opponent’s inferred SVO in degrees, we ran the learning model with each participant’s estimated parameters through the learning stimuli 1000 times per participant per social agent and calculated the resulting SVOs in degrees (see Fig. S3A). We used both the mean (μ) and the standard deviation (σ) of the distribution of calculated SVOs from these 1000 simulations as the best estimate of the opponent’s inferred SVO and the participant’s uncertainty about this estimate, and included these scalar values for each participant in our regression model (see Fig. S3B and legends).

To control for potential outliers in the population, we performed a leave-one-out cross-validation procedure and repeated the described multiple linear regression analysis. Further analysis which was done on the regression coefficients by performing 1-sample t-tests from baseline, suggested that the uncertainty term has an overarching influence on both the social risk and probability weighting parameters (all |t(49) | >5.02, all p<.001), whereas one’s own SVO uniquely contributed to one’s risk attitude only in social interactions with an individualistic opponent (t(49)=−9.66, pc.001; all Bonferroni corrected, set level for the p-value is 0.003 for 16 comparisons, see Fig.7).

**Fig. 7.**
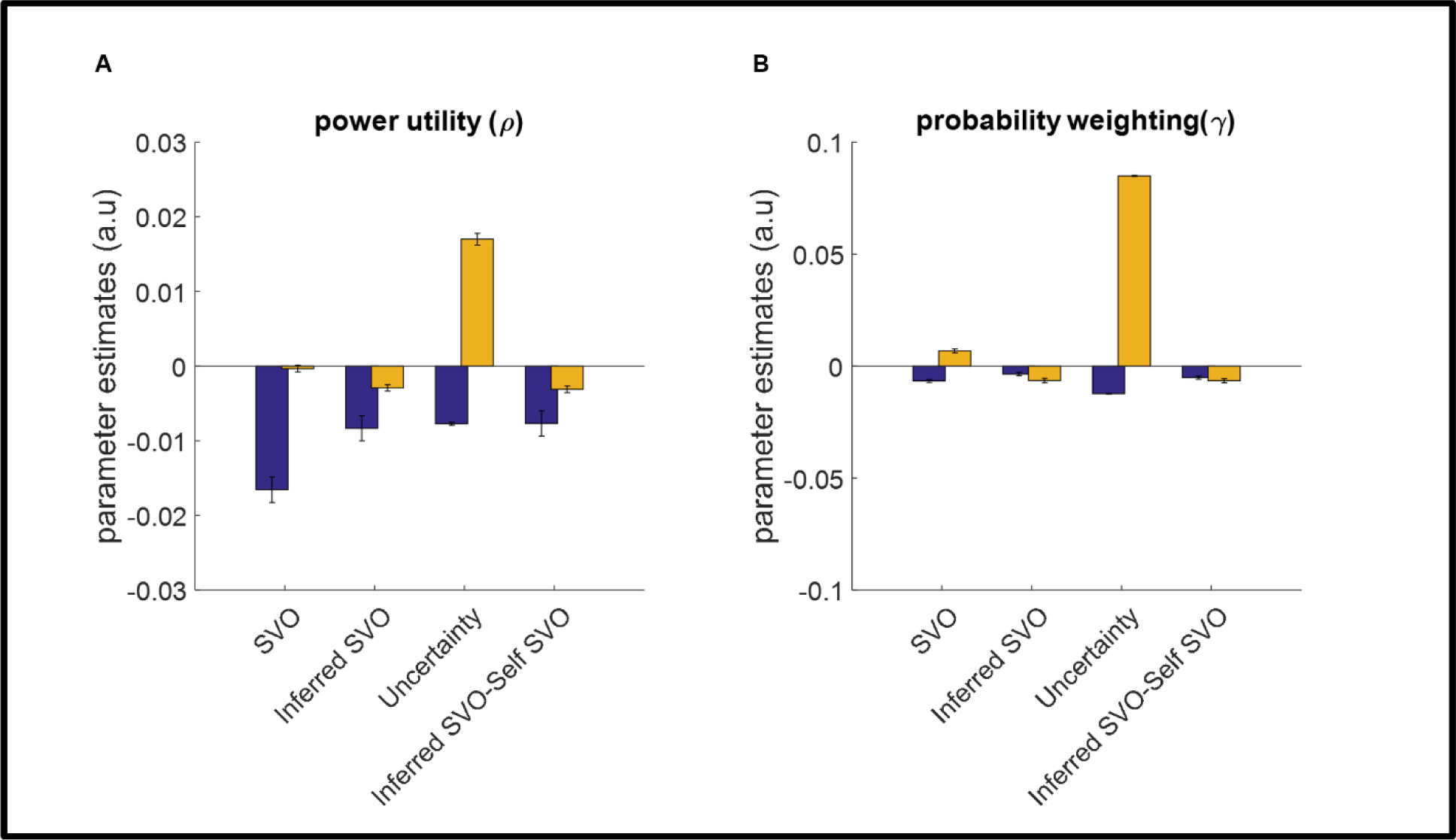
Parameter estimates {*β*_1_…*β*_4_} from two leave-one-out cross-validated multiple linear regression models accounting for the social risk and probability weighting parameters (y-axis in panels A and B, respectively). Variables in the x-axis refer to those described in the main text: SVO, participants’ Social Value Orientation; Inferred SVO, *SṼO*_*μ*_ Uncertainty, *SṼO*_*σ*_ Inferred SVO-Self SVO, (*SṼO*_*μ*_ — *SVO*). Apart from the effect of SVO on the power utility parameter for the prosocial agent (orange bar in panel A), all regression coefficients are significant (Bonferroni corrected, set level 6×10^−5^).

Finally, we wanted to provide a complementary model-free validation of the linear regression model, which showed that the variables that describe the participant’s and the opponent’s degree of prosociality influence how people adjust their risk attitudes in social interactions (see Fig 7A). If our approach is correct, participants’ frequency of choosing risky options (see Fig 4A), the risk parameters (*ρ*) estimated from the Ultimatum giving experiments, and the predictions of the model described above should line-up reasonably along the diagonal of this 3-dimensional parameter space. subsequent analysis conducted on these 3 variables suggested that the predicted risk parameters were significantly correlated with both the estimated risk parameters and the participants’ frequency of choosing risky options in the Ultimatum giving experiments (all r(49)>.383, all p<.006, Bonferroni corrected for 6 comparisons, see Fig. S4 and legends), providing converging evidence that supports of our model.

**Response Times.** Visual inspection of response time (RT) histograms suggested that RTs have a skewed distribution that violates the normality assumption. Following the recommendation of Whelan 2008 (12), we analysed the distribution of RTs across three domains (i.e. risk decision-making, social learning and social decision-making). Distributions were comparable within social learning and decision-making domains (Kolmogorov-Smirnov tests; all D≥0.12, all p>.50).

In a complementary analysis, we excluded data from 5 participants which were clear outliers. We transformed the concatenated data using a Box-Cox transformation (13) implemented in MATLAB. Overall, there was a significant main effect of the experiment type (i.e. risk decision-making, social learning and social decision-making) on RTs (F(2,222)=22.623, p<.001). RTs in the social decision-making experiments were significantly longer than those in the social-learning experiments (t(178)=6.576, p<.001), highlighting the suitability of the value-based risk model applied to the social decision-making context, where we predicted that our participants should integrate their opponents’ inferred acceptance probabilities with their self-reward magnitudes while making Ultimatum offers. The RTs were comparable both within the social learning and decision-making experiments between individualistic and prosocial agents (all t(88)<.141, all p>.164; Fig. 8).

**Fig. 8.**
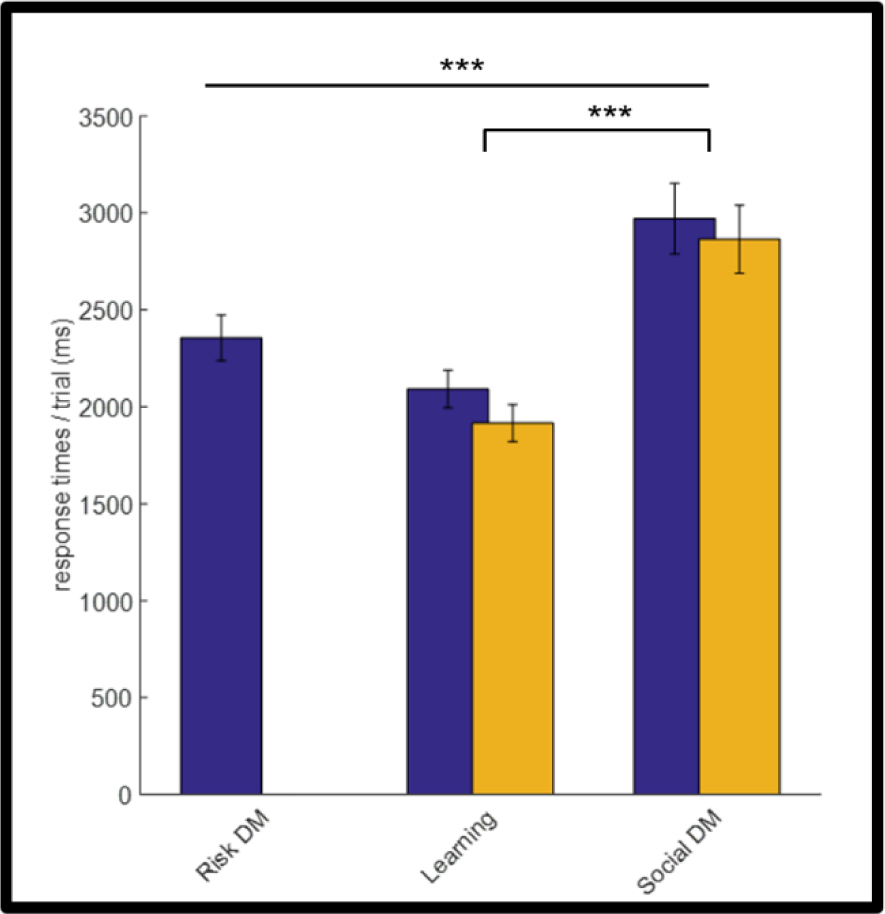
Response times across all experimental sessions. Response times between 3 domains were significantly different. Response times were longer in the social decision-making experiments relative to the social-learning experiments (*** P<0.001). Across social-learning and social decision-making experiments, there were no significant differences between prosocial/individualistic conditions.

## Discussion

In line with our prediction, the present results suggest that in social interactions which involve resource distributions between two individuals, people employ a cognitive model which shares properties with well-established computational models of value-based decision-making under uncertainty (14–16). However, unlike risk decision-making paradigms in which probability and reward information is usually given explicitly (17–20), in social interactions people need to infer others’ valuation processes (Fig. 2C) in order to make value-based decisions (21). Here, we showed that risk perception is an overarching process across non-social and social domains, whereas cognitive processes related to probability weighting are unique to social interactions in which the opponent’s valuation processes needs to be inferred. Even though social interactions demand higher-level cognitive inferences, human participants manage to make decisions nearly optimal decisions, as they would in a non-social context (Fig. 6). This near-optimal decision-making is achieved by people’s ability to adjust their decision parameters adaptively to the changing characteristics of their opponents (Fig. 7 and Fig. S4). The independence of risk and probability weighting parameter estimates within each domain are particularly important, because with a careful sequencing approach similar to a gradient decent, we were able to minimise the correlation between decision variables (22) over 2×10^6^< iterations, and make the face values of the reward distributions in the Ultimatum giving experiments identical for both opponents (i.e. prosocial/individualistic social agents, where the order of presentation was randomised for each subject and for each type of opponent). Our experimental design required participants to utilise their opponent’s encoded value function to compute the expected value difference between two options that were only cued by the colour of the icon representing their opponents (Fig. 1C). This approach minimises the possibility that within-subject differences in risk and probability weighting parameters (Fig. 4) were due to differences in the numerical components of the stimuli. In a control experiment, we further demonstrated that having an independent risk attitude is unique to situations which mimic real-life social decision-making, where people need to infer their opponents’ acceptance probabilities.

In mainstream economics and finance, people’s risk preference is often regarded as a hard-wired trait. However, a number of recent studies have suggested that risk parameters may be subjected to influence after observing others’ decisions performed in the same context(23–25), challenging this view. Here, by focusing on social interactions, we provide evidence to suggest that even in the absence of observations of comparable decisions as used by these previous studies (23–25), people’s risk-preferences may be subject to change in a social context depending on the nature of the social interaction they are engaged in. While the impact of social framing on risk-preferences has previously been investigated in the context of the Trust Game, where outcome probabilities were explicitly stated (26), to the best of our knowledge the present study is the first to describe the value computations underlying how human participants choose between Ultimatum offers where the outcome probabilities are directly related to inferences about the value functions of opponents.

It is worthwhile to emphasise that our experimental design did not have any element of observing others’ risk preferences. Instead, as we have shown, we anticipated that the perception of risk should emerge naturally due to the fact that the outcomes of these social interactions were probabilistic. We propose that “social” decision parameters may be adaptive to accommodate different interpersonal negotiation scenarios. Our subsequent multiple linear regression analyses (Fig. 7) provide evidence to support this claim, considering that key social variables related to human prosociality make differential, but mostly significant contributions to fine-tune risk (Fig. 4) and probability weighting (Fig. 5) parameters irrespective of the opponent’s SVO. Although designing two different computerised social agents increased our experimental difficulty in terms of number of the trials our participants needed to complete (t=700), it also allowed us to reveal these overarching contributions, which we think are very important for developing cognitive/computational models of social interactions.

It is necessary to comment on why we decided against the inclusion of a “competitive” agent (based on the definition of Murphy et. al) in our experimental design. Previous studies with relatively large sample sizes investigating SVO in the population showed that the population density of competitive individuals are only around 9% (27). Furthermore, people with competitive SVO are driven by achieving superiority over others, which limits them to only accept offers that satisfy this superiority criterion, making their underlying value function unsuitable for probing risk perception in social interactions. Additionally, the inclusion of a “competitive” agent would require our participants to complete at least an additional 300 trials (across social learning and decision-making sessions), making it feasibly difficult to achieve. Considering that in our cohort participants’ SVOs and participants’ inferences about the SVOs of their opponents showed a healthy degree of variability and focusing on each pairwise combination (n=100, see Fig. S5), we think our proposed model suitably meets the generalisability criteria to account for value computations in the many different social interactions that occur in real life.

Our study also has implications for understanding interactions in the Ultimatum Game, where the wide majority of the previous literature focused on responders’ behaviour (3, 28-33). In our study, the participants were explicitly instructed to treat the binary options they were presented with like thoughts in their mind, such that they knew their opponents can only see the chosen offer and can never know whether the unchosen option was better or worse. The structure of our experiment which allowed our participants to make offers from a binary selection complements previous Ultimatum Game studies where responders were commonly asked to make decisions about a single offer per trial. Under these conditions, a power utility parameter modulating self-reward magnitudes and a probability weighting parameter modulating other’s inferred acceptance probabilities described the best fitting model to participants’ choice behaviour (Fig. 3B). Participants’ self-reporting also suggested that both the self-reward magnitudes and the others’ inferred acceptance probabilities need to be considered while making Ultimatum offers (Fig. 3A). To the best of our knowledge, the current study is also the first to address value computations underlying Ultimatum giving, and it provides evidence to suggest that proposers’ do not solely rely on responders’ acceptance probabilities, but make offers based on their expected value. Based on these findings, we recommend that future cluster(34) or hyper-scanning(35) studies of Ultimatum bargaining in neuroeconomics should consider computational models which explicitly parametrize participants’ risk and probability weighting preferences.

Finally, although the social-learning session was not our main focus in this work, we showed behavioural computational evidence to suggest that our participants could suitably transfer the encoded value functions of others from one (learning) environment (i.e. observing social agents’ responses to singular Ultimatum offers) to another in which they were asked to solve an optimal social decision problem (i.e. making Ultimatum offers from binary options). We showed evidence to suggest that human learners do not represent their opponents in terms of large numerical self and other reward grids to encode their value functions like a Bayesian Ideal Observer model would do. As a result, in the context of our current experiment, a simpler social value orientation model predicted human social learning behaviour better. These results are mostly in line with previous learning literature which put forward Bayesian Ideal Observer models to reveal hidden parameters of a generative process (e.g. estimated outcome volatility; Behrens et al., 2007, Browning et al., 2015, Pulcu and Browning, 2017) but do not necessarily predict participant choice behaviour better than simpler learning models. We think that our behavioural study highlights the need for further research in three main streams, ideally involving functional magnetic resonance imaging (fMRI): (i) what are the regions involved with neural computations underlying how people transfer the encoded value functions of their opponents in social interactions; (ii) what are the neural mechanisms responsible for tracking the value functions of opponents with different SVOs; and (iii) which brain regions encode the estimated trial-by-trial variability in the social risk and probability weighting parameters.

## Author Contributions

E. Pulcu and M. Haruno developed the study concept and contributed to the study design. Testing, data collection, data analysis and interpretation were performed by E. Pulcu under the supervision of M. Haruno. E. Pulcu drafted the manuscript, and M. Haruno provided critical revisions. Both authors approved the final version of the manuscript for submission.

## Acknowledgements

The study was funded by CREST, JST, COI to Osaka University, JST and KAKENNHI (17H06314 and 26242087) awarded to M. Haruno. The funders had no role in study design, data collection and analysis, decision to publish or preparation of the manuscript. We are grateful to Satoshi Tada for technical assistance and Dr. Michael Browning, Prof. Jon Roiser and Prof. Mauricio Delgado for their recommendations on an earlier version of this manuscript.

## Materials and Methods

**Participants.** In total 50 healthy individuals (54% males) who reported no history or current neurological or psychiatric disorders, or use of any psychotropic medication were recruited from the general population. The average age of this cohort was 31.5 (range: 20-56 years; STD=±9.49). On average the participants had 16 years of education (STD= 2.1 years) and reported annual income of 1.91 million Japanese Yen (STD=1.71 million Yen). This cohort was recruited from the general population with a convenience sampling approach and contained a higher percentage of prosocial individuals (n=29 vs n=5 individualistic participants according to the categorical classification of the SVO Triple Dominance Measure(36) which penalises inconsistent responses).

**Experimental Procedures.** The study took place at the Center for Information and Neural Networks (CiNet) and was approved by the CiNet Research Ethics Committee. Participants who met the inclusion criteria were given an appointment for the behavioural experiments. The testing session began by an explanation of the research procedures, followed by obtaining an informed consent. Prior to any experiments, the participants completed a battery of questionnaires related to their demographic information and Social Value Orientation (SVO) in pen and paper format.

Before the computerised experiments, the colour coding of icons was explained to the participants (see Fig 1). The red icon always represented the participant him/herself. In the paradigms which had a social component, the blue icon always designated the individualistic opponent and the orange icon always represented the prosocial opponent. Because the computerised testing involved completing 700 trials across different paradigms, we wanted to minimise participant errors (e.g. due to fatigue) by keeping the colour coding consistent throughout the testing session.

The participants first completed the value-based risk decision-making task which lasted for 100 trials (Fig. 1A). This task was designed to capture participants non-social value-based decision-making preferences at baseline. The risk decision-making task involved binary decisions between two probabilistic gambles, where outcome reward magnitudes (between 10 and 100 Japanese Yen) and outcome probabilities were generated by MATLAB’s *rand* and *randsample* functions (for probabilities and reward magnitudes, respectively). These decision variables were shuffled until they were decorrelated, and the expected value difference between the options, calculated based on Eq. 5 and 10 (see below), was normally distributed with mean ~0. These features of the stimuli allow the fitted stochastic choice function to capture how participants behaviour shift from one option to the other in relation to the changing expected value difference between options, from negative to positive.

After this stage, the participants completed the learning sessions for both the individualistic and prosocial agents, in which they were asked to predict whether the social agents will accept or reject the offers coming from other anonymous individuals. The Ultimatum offers were also generated to be between 10 and 100 Japanese Yen. The participants were told that the social agents whose Ultimatum responses they needed to predict were two participants from a previous study conducted by our research group. In fact, they were computerised agents making decisions following an underlying value function (see below for details). A similar methodology is frequently used in behavioural studies conducted by other research groups (25, 37). The learning sessions contained 180 trials each, and the order of the learning sessions was counterbalanced across participants. The participants won ¥25 for every correct prediction, which was added to their performance-based reimbursement. Incorrect predictions did not change participants’ running total. After completing the learning sessions, participants were asked to respond to various descriptive questions while considering the imagined personalities of these social agents (see the full list of questions in Fig. S1 legend).

Finally, the participants completed the social decision-making experiments, for 120 trials against each social agent in the same order they completed the learning sessions. With careful sequencing of Ultimatum offers presented in the social decision-making stage, we were able to make the face values of the binary Ultimatum offers identical for each social agent. Consequently, the participants needed to use the value function of different opponents to compute the expected value difference between the binary offers in the Ultimatum giving experiment, even though their face values were the same. This approach allowed us to control for any change in decision parameters which can be attributed to the numerical differences in the stimuli. The presentation order of the trials was purely randomised across all participants. In the Ultimatum giving experiment, participants obtained the monetary amount (R_0_) in all of their accepted offers. This amount was also added to their performance-based reimbursement. All of the behavioural experiments were self-paced, and participants were paid the total amount they accumulated across social learning and all decision-making experiments (both non-social and social experiments; i.e. all participant decisions had real-life financial consequences). All experiments took place in a comfortable room designated for testing purposes and all tasks were presented by PsychToolbox 3.0 running on MATLAB (MathWorks, Inc.).

**Description of the computerised social agents.** Two distinct computerised social agents, whose behaviours were guided by the way they computed the subjective value of the Ultimatum offers they faced were defined by a model from the social value orientation (SVO) framework based on a model derived from a previous publication by our research group (1). Here, the subjective value (V) of *a* condition is calculated as:

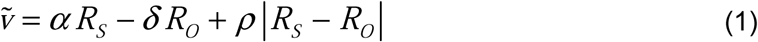

where *R*_*S*_ (always from the perspective of the social agents) and *R*_*o*_ depicts self and other’s reward magnitude, respectively. The agents make decisions following a stochastic choice model where *q*_*A*_ the probability of accepting a condition (38):

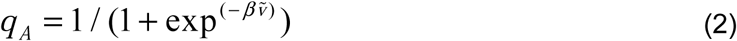

Here, *β* is the inverse temperature term, which gives the shape of the sigmoid function based on ouro *priori* assumption about the shape of the sigmoid function in humans, which should be the case if the number of trials in the learning session approaches to infinity; that is, if the participants had the opportunity to observe the behaviour of the computerised social agents for a very long time.

The hyper-parameters defining the valuation of the agents, *α,ε, ρ, β* were set to [1.096, 0.382,-2.512, 0.037] for the individualistic agent; and [1.368,-0.644,-3.798, 0.045] for the prosocial agent. The key difference between these two agents was that the prosocial agent valued conditions cooperatively, whereas the individualistic agent valued them competitively, and that the prosocial agent was more sensitive to the absolute value difference between the self and other’s reward magnitude. We generated a vector of responses to 180 trials in the learning sessions by the defined model, where the SVO of the social agents was calculated by the following formula derived from Murphy et al. 2011:

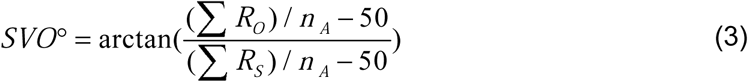

where *η*_*A*_ is the total number of accepted conditions. After extensive simulations to evaluate how agents would behave, a selection was made such that the SVO of the prosocial agent was 31.47° and the SVO of the individualistic agent was 12.36°; clearly falling into the categorical boundaries described by Murphy et al., 2011. Therefore, these strategies were labelled as “prosocial” and “individualistic” throughout this manuscript.

In order to make sure that the choice behaviour of these computerised agents will adequately mimic decisions of real human participants, we conducted an additional control experiment in which participants in an independent cohort (n=40; age: 21±2.1; 60% males; SVO: 25.7±15.6) responded to Ultimatum offers coming from different proposers, as it was in the social learning sessions of the main experiment. We simulated the choices of the computerised agents 100 times for each participant and investigated the extent to which their decisions coincided with the decisions made by real humans. This control experiment suggested that the behaviour of computerised agents would be well tolerated in the main experiment, particularly considering that the offers being evaluated covered the indifference point (i.e. expected value difference near 0), which is where the choices of the computerised agents and real human participants would be random (~50% accept, see Fig. S6 and legends). In addition to this numerical analysis conducted in an independent cohort, it is important to point out that our participants were able to identify at least ~4 people in their close circles whose decisions resembled the decisions made by the computerised agents (see Fig. S1, responses to Q3), suggesting that the behaviour of the computerised agents in the main experiment was overall well tolerated.

**Social-learning.** There are a few models in the literature that account for how people learn during social interactions (21, 39, 40). Due to the widely-known complexity of this process (41, 42), we did not focus too much on specific models of social-learning by performing detailed trial-by-trial analyses here. Previous studies suggested that using the Ultimatum Game as an environment to investigate how people learn other’s social preferences is challenging due to the fact that the game structure has a strong non-monotonicity (42), as people are shown to be sensitive to unfair resource distributions irrespective of whether they are favourable or not. Nevertheless, the social learning experiments were still necessary in order to model how inferred probabilities are processed during the Ultimatum giving experiments, in which we anticipated that our experimental setting will naturally reveal how our participants evaluate the risks associated with the possibility of others rejecting their offers. Therefore, we modelled this first step based on the assumption that learning occurs through the successful simulation of another’s valuation model (43) (achieved by model-free, reinforcement or Bayesian learning, or their weighted combination) such that as the number of trials (*t*) approach to infinity, 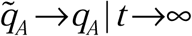, that is, the participant’s inferred acceptance probability 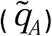 will fully converge to the social agent’s true acceptance probability (*q_A_).* We confirmed that social learning occurs suitably well by performing model-free analysis of the data from the learning sessions and also by fitting the proposed valuation model of the social agents to the participants’ choice data to generate the participants’ inferred choice probability for each social agent (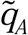, see Fig. 2).

Although the learning session was not our primary focus, we still wanted to compare the performance of the social valuation model with two alternative Rescorla-Wagner models (44) and one Bayesian Ideal observer model, which were fitted to the participants’ choice behaviour in the learning sessions.

In alternative Model 1, the participant updates his/her estimate of the social agent’s overall acceptance probability on trial *t* in proportion to the prediction errors (ε) on trial *t-1* on a trial-by-trial basis:

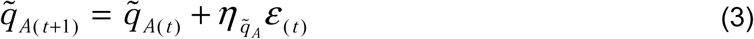

where 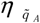 is the learning rate. In alternative Model 2, each of the social agent’s parameters in the described SVO model (Eq.l and 2) is updated on a trial-by-trial basis, for example:

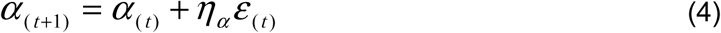

where *η_α_* the learning rate updating the value of the free-parameter *α* from trial t to *t*+1. This second model has 4 free parameters that represent the learning rates by which participants updated each of the parameters of the SVO-based valuation model (Eqs. 1 and 2).

The Bayesian Ideal Observer model would represent each Ultimatum offer over a numerical grid of reward magnitudes for the self and the other. The model would start with flat priors (i.e. all inferred acceptance probabilities set to 0.5 on trial 1; a=1, b=1) and learn the other’s Ultimatum preferences by updating the mean of the nested beta distributions 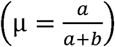 over this numerical grid following each observation (e.g. a=a+1 after each accept and b=b+1 after each reject response). Because the 180 trials that our participants completed in the learning sessions are not enough to cover the whole numerical grid, we used a 3×3 smoothing kernel over the model’s inferred acceptance probabilities (see Supplementary Video for the behaviour of the Bayesian model throughout the learning session), allowing the model to make inferences dynamically for seemingly comparable offers as humans would do.

Model comparisons for the learning session favoured the SVO-based valuation model, which had significantly lower-log likelihood values relative to the reinforcement learning models (all F_2,147_>168, all p<0.001, Bonferroni corrected). Furthermore, the close relationship observed between *q*_*A*_ and 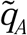 from the SVO model as well as model fitting metrics falling into the highly desirable range (11) suggest that the present approach is suitable for obtaining the participants’ inferred acceptance probabilities. Similar approximations have been used by other research groups to reduce the model complexity of learning in social decision-making tasks (21).

**Description of the computational models for the control experiment.** Considering that our social agents were designed to make choices following specified value functions, interacting with them in the main experiment should naturally probe a perception of risk. Consequently, we decided to select a value-based risk decision-making task (Fig. 1A) as our control experiment. This selection enabled us to understand whether people use an overarching computational model in non-social and social contexts for value-based decision-making under risk and uncertainty. We fitted various computational models to the participants’ choice behaviour, as described below.

In line with the previous literature, we modelled value-based decision-making in a way which allows human participants to make binary choices between probabilistic gambles by computing the expected value (π) of the options they face:

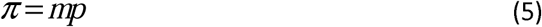

where *m* is the reward magnitude, *p* is the probability associated with an option, and ρ is their multiplicative integration (Model 1). However, previous work showed evidence for nonlinear modulation of outcome probabilities in human value-based decision-making (25)(45). In order to capture these processes, we utilised a probability weighting function based on previous studies (43, 46):

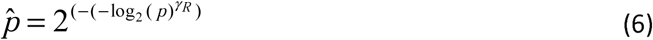

where *γ_R_*>0 is a free parameter which modulates actual outcome probabilities nonlinearly into subjective probabilities. The log_2_ function always crosses the p/p diagonal at 0.5 and in our point of view accurately captures the intuition that people should have a somewhat accurate perception of 50/50 odds. Participants then compute the expected value of a gamble accordingly (Model 2):

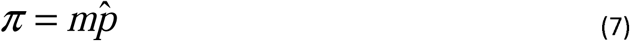

In an alternative model (Model 3), we considered an expected value computation which can account for participants’ risk-preferences by revealing the curvature of their utility functions:

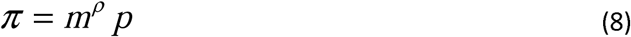

where *ρ* is the power utility parameter, with *ρ >* 1 indicating a risk-seeking, *ρ <* 1 indicating a risk-averse and *ρ*= 1 indicating a risk-neutral preference.

Finally, the full model (Model 4) considered both of these nonlinear processes in computing the expected value of an option:

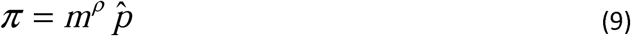

Across all models, it is assumed that participants make their choices in relation to the subjective value difference between each gamble (i.e. the difference between the left and right options):

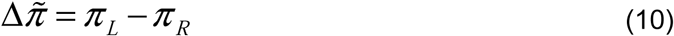

and trial-wise stochastic choice probabilities for each gamble are generated by a sigmoid function:

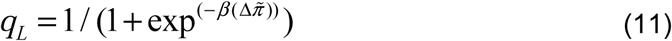

where *β* > 0 is the inverse temperature term adopted from thermodynamics and determines the degree of stochasticity in participants’ choices.

**Description of the social decision-making models.** In the main experiment, the participants were asked to make Ultimatum offers to the social agents. If their offer was accepted, participants would receive the amount *R*_*O*_, whereas if their offer is rejected both sides got nothing for that trial (as in a typical Ultimatum Game experiment).

It is possible that participants completed the social decision-making task using a number of different strategies. Here, we considered cognitive models with variable complexity, where our preferred model proposed that participants simulate the social agents’ acceptance probability for both the chosen and the unchosen options and compute the expected value difference between the options by integrating these inferred acceptance probabilities with their self-reward magnitudes. We formally define these different models below.

According to Model 5, the participant’s decision value (Δ*ṽ*) depends on the difference between the social agent’s inferred choice probability 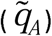 that is associated with the offers on each side {*L, R*}, whereby the participant makes decisions only by considering the opponent’s acceptance probability (i.e. choosing the offer which they think is more likely to get accepted):

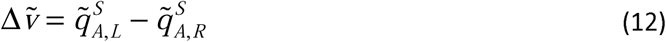

where *S* ϵ{*i, p*} defines whether the opponent is individualistic or prosocial.

Due to the probabilistic nature of the social decision-making task, we hypothesised that the inferred acceptance probabilities could also under go a similar probability weighting transformation as in the non social value-based decision-making task (Eq. 6). Therefore, in Model 6, we considered that participants’ decision value (Δ*c*) may be computed by the following two equations:

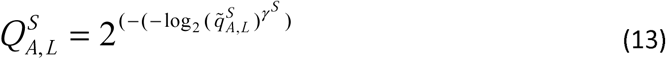

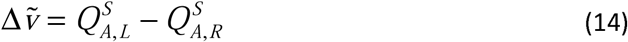

where *γ^s^*>0 is a free parameter accounting for the nonlinear modulation of acceptance probabilities in the social decision-making task, which are estimated separately for each social agent. Here, the introduction of a free parameter is critical and allows us to evaluate differences between the modulation of probability weighting in social and non-social contexts.

In Model 7, we considered a more sophisticated value computation that also integrates participants’ own payoff (*R_O_*). This computation assumes that participants employ a cognitive model with a large degree of overlap with the previously defined non-social value-based decision-making model (i.e. Model 2), whereby the participant can choose to make an offer with lower 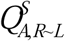 if the overall expected value is higher.

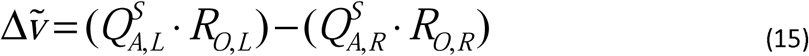

In Model 8, which largely overlaps with the non-social value-based Model 3, the decision value is computed by the following formula which would account for the participants’ risk preferences by revealing the curvature of their utility functions (25):

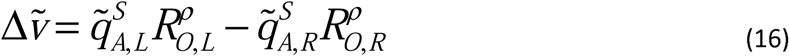

where *ρ* is the power utility parameter, with *ρ* >1 indicating a risk-seeking, *ρ* <1 indicating a riskaverse and *ρ* = 1 indicating a risk-neutral preference.

In Model 9, which largely overlaps with the full non-social value-based Model 4, the decision value is computed by the following formula which accounts for both the participants’ risk and nonlinear probability weighting preferences:

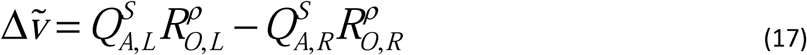

After each social decision-making experiment, we asked participants to rate how much they considered other’s acceptance probability, their own payoff, or both on a 0-to-10 scale (e.g. a rating of 5 meaning integration with equal weighting; see Fig 3A for self-reported integration weights). In Model 10, we used this rating as a linear weighting information, where the weight parameter, *W*, takes a value in the normalised space (i.e. between 0 to 2) with 0.2 increments because it was directly derived from the participants’ own report on a 0-to-10 scale (e.g.*W*=1 indicates integration with equal weighting,*W*=0.4 the indicates more weight is given to the self-reward magnitude, etc.). In essence, this model is similar to non-social value-based Model 1, but with added subjective integration weights. Here, the decision value is computed as follows:

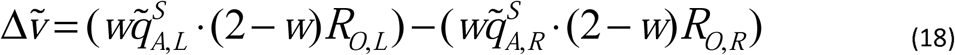

Finally, as a baseline control condition, we also investigated the fitness of a model which makes decisions based on self-value difference alone (i.e. Model 11; 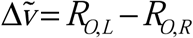).

At the very last step, choice probabilities under each model *M ϵ* {5,6,7,8,9,10,11} were generated by a sigmoid function:

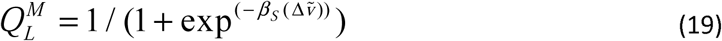

where *β_S_*>0 is the inverse temperature term estimated separately for the social decision-making experiments.

We used a Maximum Likelihood Estimation procedure to evaluate how well the proposed cognitive models explained our participants’ choice behaviour. The free parameters were estimated using a non linear optimization method over a numerical grid which covered the whole parameter space, using MATLAB’s (MathWorks, Inc.) *fmincon* function with random starts.

We selected between competing models based on their Bayesian posterior probability weights given the data (47) by the following formula:

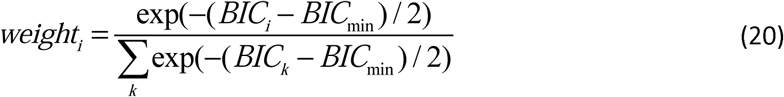

while considering each model’s Bayesian Information Criterion (BIC) value (48), which penalises more complex models with additional free parameters.

For robustness, we also implemented a complementary Bayesian model selection approach (49, 50) by feeding a matrix (number of participants X number of competing models) of log likelihood values for each model to the readily available scripts from the SPM12 library (spm_BMS; www.fil.ion.ac.uk/spm) and computed the exceedance probabilities for each of the competing models.

## Supplementary Figures and Legends

**Fig. S1.**
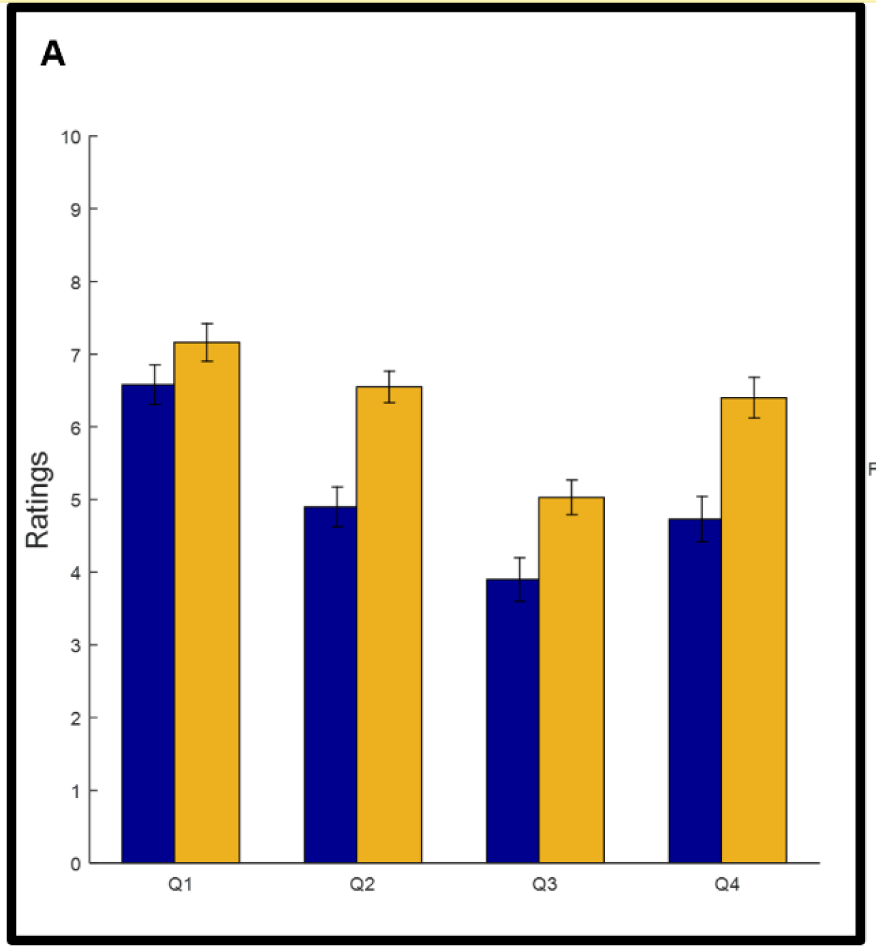
Descriptive ratings participants made about social agents following the observational social-learning sessions. (A) Participants responded to a number of questions while thinking about the personality of these social agents (all rated on a Likert scale from 0 to 10). Q1: How much do you think this person cares about rewards to others? Q2: How much would you like this person if you spent 1 hour with him/her in real life? Q3: How many people do you know in real life who resemble this person? Q4: How socially close do you feel towards those people that you know? A fitted 4×2 MANOVA suggests there is a significant main effect of the social agent (F=3.01, P=0.03) and a significant effect of the interaction term (F=10.16, P<0.001).

**Fig. S2.**
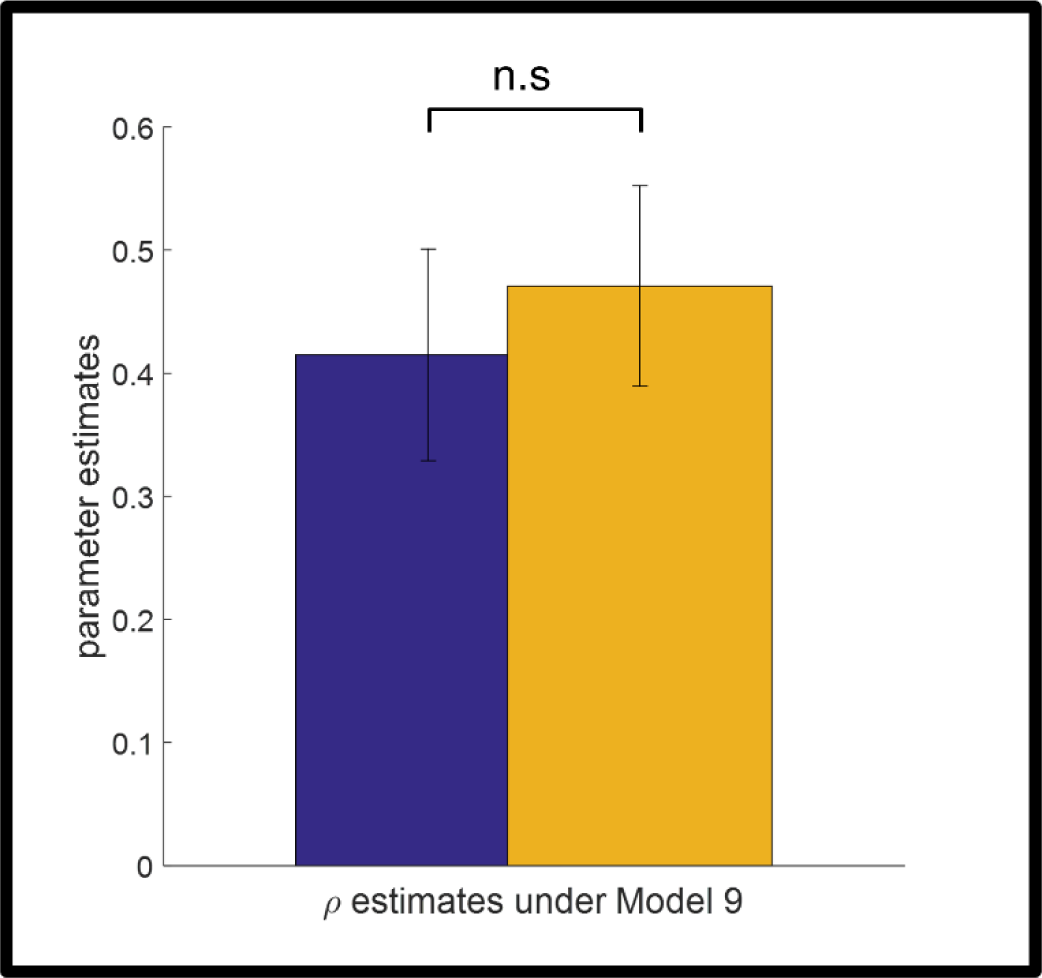
Risk parameters 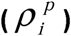 estimated from the Ultimatum giving experiments against two different social agents with different SVOs were not significantly different (P=0.64, n.s: not significant).

**Fig. S3.**
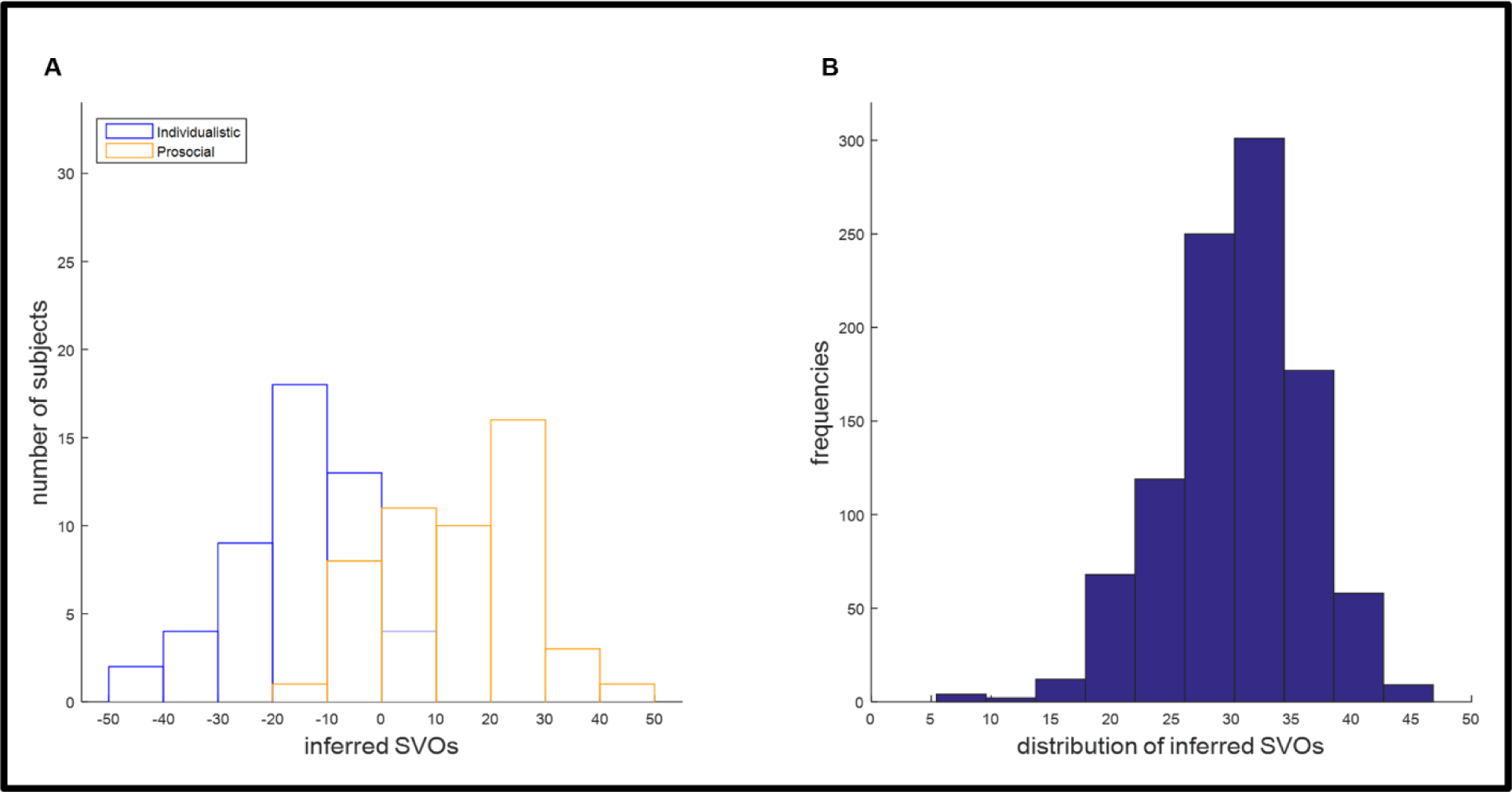
Distributions of opponents’ inferred SVOs (*SṼO*) based on simulations of the encoded value functions 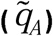 from the social-learning sessions. (A) The distribution of inferred SVOs for the whole cohort (i.e. distribution of means obtained from each subject’s simulation) based on 1000 simulations of the encoded value functions per participant per social agent (i.e. prosocial and individualistic). These distributions were significantly different from each other (*D*=0.78, P<0.001), suggesting that participants were able to make distinct inferences about the SVOs of their opponents. (B) The distribution of inferred SVOs in 1000 simulations from a single subject gives a normal distribution of inferred SVOs, of which the mean and the standard deviation were used as input variables for the multiple linear regression analysis. Here, the true SVO of the prosocial agent was 31.5°, which falls close to the mean of this distribution (*μ* =30.77° ± 5.4).

**Fig. S4.**
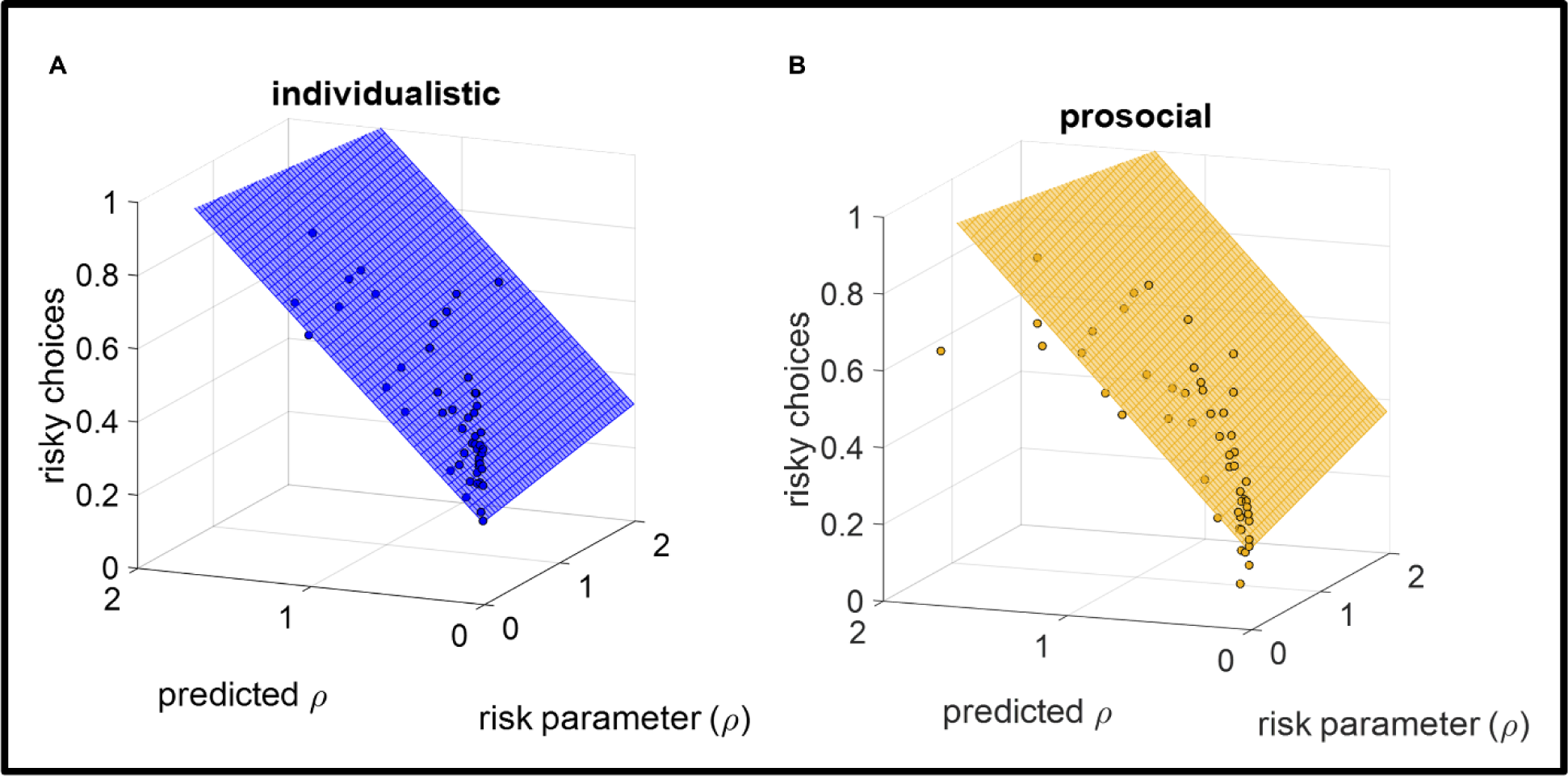
A 3D scatter plot with fitted planes through the least squares regression lines, summarises the relationship between the participants’ normalised frequency of choosing risky options (i.e. the vertical axes in both A and B; identical to Fig. 4A), the estimated risk parameters in the Ultimatum giving experiments and the predictions of the multiple linear regression model. Predicted values of the risk parameter (*ρ*) in the Ultimatum giving experiments correlated significantly with the actual parameter estimates and the participants’ choice frequency (r>.383, p<.006, Bonferroni corrected).

**Fig. S5.**
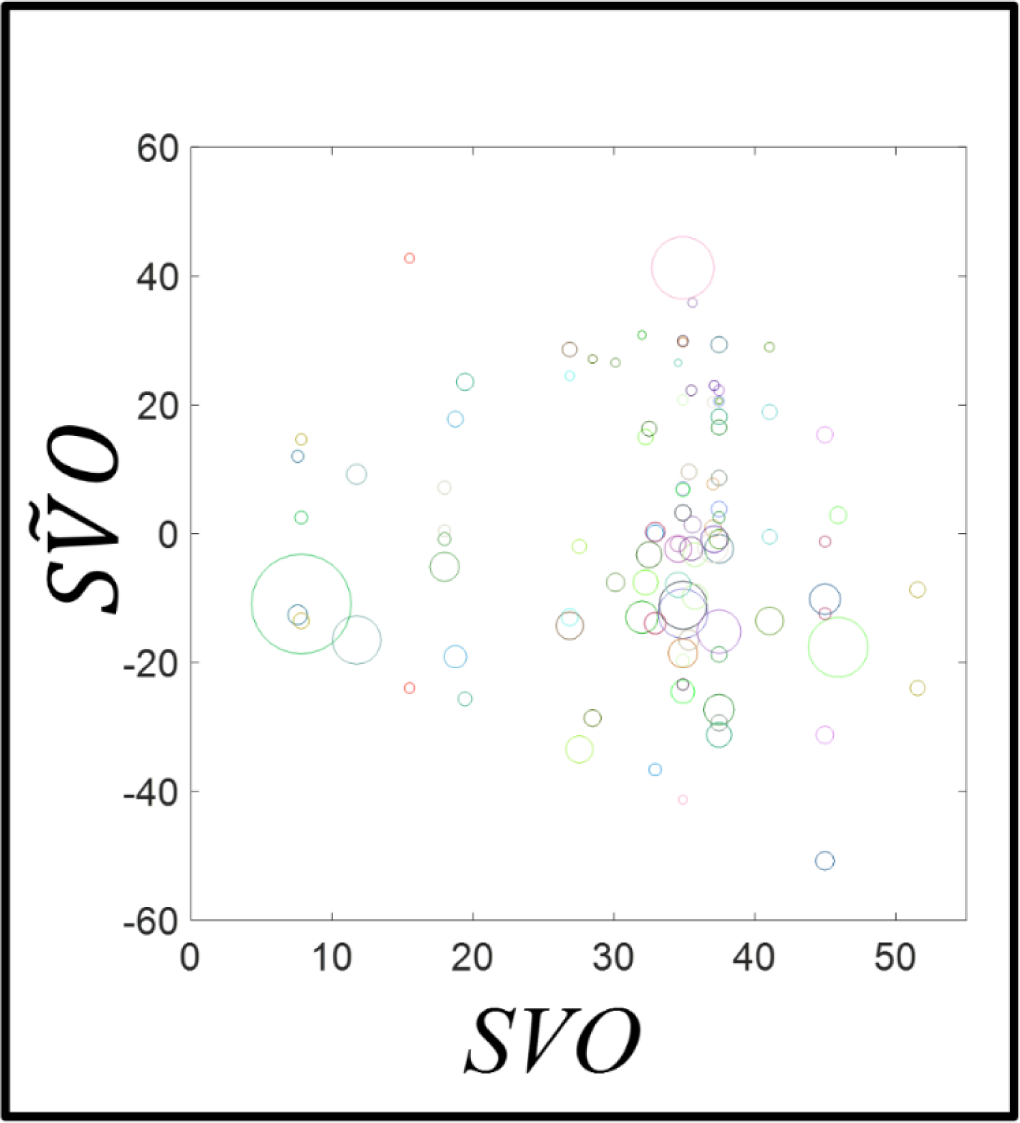
Combinations of social interactions covered by the current study (n=100) with respect to the participants’ own (x-axis) and their inference of their opponents’ SVO (y-axis). Markers with the same [R, G, B] colour coding refer to a single subject’s data point. Marker sizes are proportional to the uncertainty estimates (*SṼO*_*σ*_, which was included as an input variable in the multiple linear regression model described in the main text.

**Fig S6.**
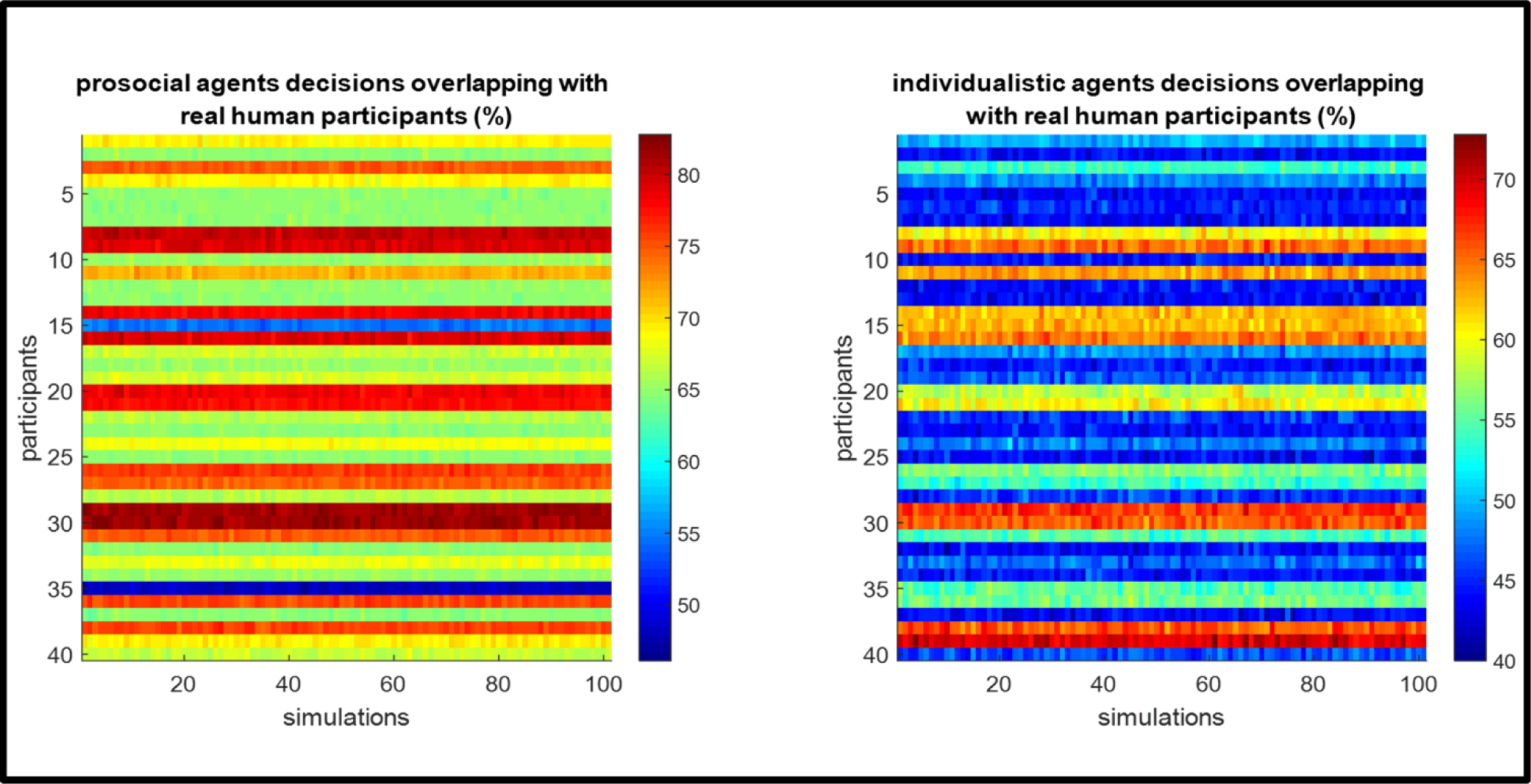
Summary results of a control experiment conducted in an independent cohort (n=40). Colour bar shows the percentage of choices overlapping between the prosocial agent and human participants responding to Ultimatum offers (left panel), and the individualistic agent and human participants (right panel). The decisions of the computerised agents were simulated 100 times for each participant, and the percentage overlap was calculated accordingly. Considering that expected value difference (~0 range) was covered with an adequate number of trials across all experiments reported in this manuscript, where the choice behaviour is stochastic, these results provide additional support to the subjective ratings reported in Fig. S3 (responses to Q3) that indicate the behaviour of the computerised agents should be well tolerated by the participants in the main experiment. https://youtu.be/pMC9H9vV9Fs Bayesian Ideal Observer model can track social agents’ Ultimatum acceptance probabilities optimally by updating the estimated means of the nested beta distributions over a numerical grid of self and other’s reward magnitudes.

